# Targets of the human antibody response to the variant surface glycoprotein of *Trypanosoma brucei*

**DOI:** 10.1101/2025.09.25.678491

**Authors:** Jaime So, Bailin Zhang, Lulu Singer, Alexander K. Beaver, Johan P. Zeelen, Philippe Büscher, Veerle Lejon, Monica R. Mugnier

## Abstract

Human African Trypanosomiasis (HAT) is a severe disease endemic to Sub-Saharan Africa caused by the parasite *Trypanosoma brucei*. Diagnosis of the chronic form of HAT relies heavily on serological screening tests that detect specific antibodies. *Trypanosoma brucei* parasites rely on the antigenic variation of their variant surface glycoproteins (VSG) to evade host antibody recognition. Although VSG is the primary target of host antibodies, studies of the antibody response to VSG have been limited by low-throughput approaches that allow for the analysis of only a few VSG proteins at a time, primarily in animal models. Here, we use phage immunoprecipitation sequencing (PhIP-seq) to characterize the antibody response to VSG in HAT patient serum. Using a comprehensive T7 bacteriophage display library containing peptides representing 12,108 unique *T. brucei* VSGs, our analysis revealed significant differences in the anti-VSG antibody between the acute and chronic forms of HAT. Our results show that, in chronic HAT, the top lobe of the VSG is the primary interface of antibody interactions. By characterizing the targets of anti-VSG immunity, we also identified a potential bias in the CATT, the most common test used for HAT screening. Finally, we were able to identify sets of seroprevalent VSG peptides for each form of the disease that successfully distinguish between infected and uninfected individuals. Our results demonstrate the utility of characterizing host antibody responses against African trypanosomes, which can reveal unique features of parasite biology while also uncovering new diagnostic targets.

## Introduction

Pathogens and their hosts are pitted against each other in a relentless evolutionary arms race. With the ability to generate specific antibodies, B cells can react to a potentially unlimited number of protein antigens. In turn, many human pathogens have evolved highly sophisticated strategies to continuously evade recognition by this flexible host antibody response. *Plasmodium*, *Giardia*, *Babesia*, *Borrelia*, and others utilize diverse surface proteins to escape host immunity by expressing one antigen at a time from a silent repertoire of variants stored within the genome^1^, a process referred to as antigenic variation.

The parasitic protozoan African trypanosomes, *Trypanosoma brucei spp.*, provide an excellent model for studying antigenic variation. The surface of each *T. brucei* cell is completely enveloped by ∼10 million identical copies of a variant surface glycoprotein (VSG)^2^. Parasites remain extracellular throughout infection, responding to perpetual attacks by host antibodies by periodically “switching” expression of their VSG coat to a different variant. Specific antibodies remain the host’s primary defense against *T. brucei*. At low antibody titers, hydrodynamic forces from parasite swimming traffics VSG-bound antibodies to the flagellar pocket where they are endocytosed and degraded^4^. This defense is eventually overwhelmed as higher titers of VSG-specific antibodies mediate parasite clearance from the bloodstream by phagocytosis or complement activation^5–7^. At this point, parasite cells must circumvent antibody recognition by switching expression of their VSG coats to antigenically distinct variants^7,8^. *T. brucei* alters its antigenic profile not only by encoding several hundred VSG genes, but also through its ability to generate new “mosaic” variants through recombination, utilizing the thousands of VSG sequences encoded within its genomic repertoire.^8^

Although antibodies mediate parasite clearance, experimental data regarding the anti-VSG response is relatively limited. Indeed, much of our understanding is inferred indirectly from structural and genomic data. The VSG protein contains two domains: a large and variable N-terminal domain (NTD) that sits above a more conserved C-terminal domain (CTD) proximal to the plasma membrane^2,9^. The NTD is considered the main interface of antibody interactions as it harbors the most antigenic diversity, typically only sharing ∼10 – 30% identity between variants^10,11^, while the less variable CTD is thought to be hidden on live parasites and thus less important for parasite clearance^12^. Packing of VSGs upon the parasite cell surface leaves the NTD of the protein exposed^9,12^, and monoclonal antibodies and nanobodies against VSG which are able disrupt live parasites bind to sites throughout the NTD^13,14^. However, direct measurements of antibody responses against VSG have typically evaluated only a few VSGs at a time, utilizing laboratory strains of the animal infective subspecies *T. brucei brucei* or purified/recombinant protein domains. Given the enormous size of the VSG repertoire, it is unclear whether the findings from a relatively small number of VSG-antibody pairs can truly be generalized to the parasite’s vast antigen repertoire.

The antibody response to VSG in the context of human infection is even more poorly understood, as most work has been performed in mouse models^15^. Indeed, to our knowledge, there has been only one study that directly analyzed antibody responses in human serum, and its focus was only on two VSGs, LiTat1.3 and LiTat1.5^16^. However, the VSG-antibody interface is critical to the serological tests used to screen for infection. Two subspecies of African trypanosome have evolved the capacity to infect humans: *T. brucei gambiense* and *T. brucei rhodesiense*. Among all reported cases, HAT caused by infection with *T. b. gambiense* (gHAT) accounts for 94% of cases and is mainly distributed throughout West and Central Africa, while the remaining 6% of cases are due to *T. b. rhodesiense* (rHAT) and are localized to East Africa^17^. While recently developed RDTs use combinations of antigen types, the longstanding gold standard method for gHAT case detection is the Card Agglutination Test for Trypanosomiasis (CATT/*T. b. gambiense*) which detects serum antibody reactivity against a single parasite strain expressing the VSG LiTat1.3^18–20^. There is no operational serological test to screen for rHAT which instead relies on case detection by blood microscopy^20^, though the WHO has expressed a need for a rapid serological test for rHAT^21^. Given the importance of serological screening, particularly for diagnosis of gHAT, a better understanding of how hosts respond to *T. brucei* infections could not only provide insight into the mechanisms of antigenic variation broadly, but also inform new strategies for HAT case detection.

In the current study, we present a comprehensive analysis of antibody reactivity against VSGs in human serum using phage display immunoprecipitation sequencing (PhIP-seq). PhIP-seq allows for sensitive measurement of antibody binding to thousands of peptide antigens simultaneously and has been successfully applied to investigations of autoimmunity, allergy, and viral infection^22–25^. Using this noninvasive approach, which requires only 1µl of serum for screening, we characterized anti-VSG antibody reactivities in hundreds of HAT patient serum samples along with matched negative controls. Our analysis revealed the primary targets of anti-VSG antibodies, while also exposing a potential bias in the CATT test used for gHAT diagnosis. Using this comprehensive dataset, we were additionally able to identify seroprevalent peptides that could serve as the basis for new HAT diagnostics. By screening antibody binding against thousands of VSGs in hundreds of patients, this study begins to capture the elusive VSG-antibody interface.

## Results

### PhIP-seq is a multiplexed assay for evaluating antibody responses against VSG

In PhIP-seq, a phage display library is exposed to a serum sample, allowing antibodies to bind the antigens displayed on the phage capsid. Phage-antibody complexes are then immunoprecipitated and submitted to targeted high-throughput DNA sequencing of phage inserts to identify reactive antigens, allowing for high resolution characterization of antibody specificities (Fig. 1A). To understand the anti-VSG antibody response, we designed a T7 phage display library containing 76,601 unique sequence inserts encoding 90 amino acid peptides tiled with 45 amino acid overlaps and representing 12,108 different VSG genes, pseudogenes, and gene fragments. VSG sequences for the three major subspecies of *T. brucei* (*T. b. brucei*, *T. b. rhodesiense*, and *T. b. gambiense*) were included along with *T. equiperdum* and *T. evansi* which are closely related to *T. brucei* (Fig. 1B)^26^. VSG N-terminal domains can be categorized into two broad types, A and B, based on shared topology and structural architecture^10^; our library design contained these types at roughly equal proportions (Fig 1B)^10,27^.

**Fig. 1).**
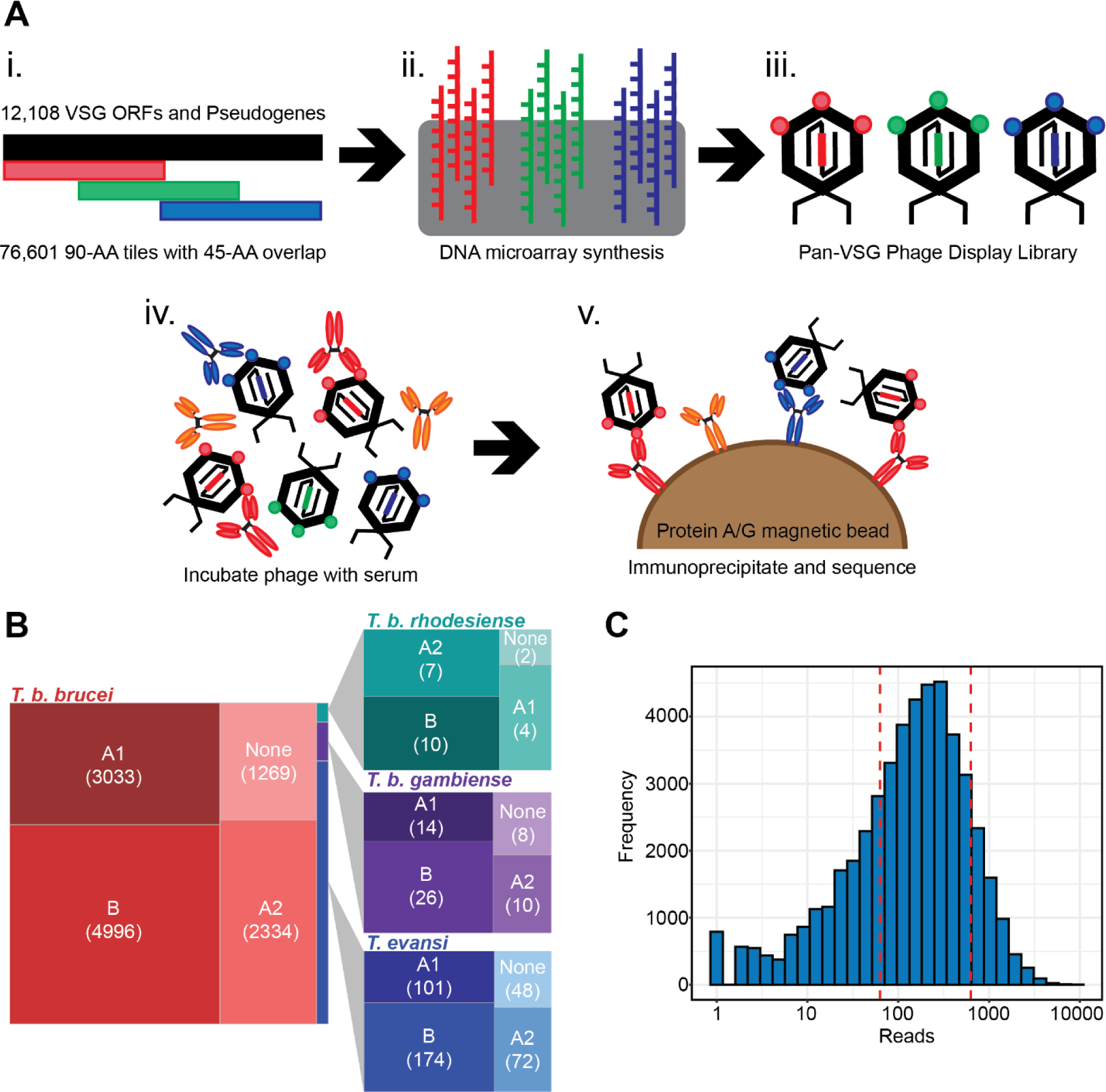
Library Design and PhIP-Seq Methodology. **(A)** (i) The VSG phage display library contains 90-AA peptide tiles that overlap by 45-AA and includes sequence from 12,108 VSG genes and pseudogenes. (ii) 300-nt DNA oligos encoding each peptide were synthesized by microarray and (iii) cloned into T7 phage display vectors. (iv) The phage library is mixed and incubated with serum. (v) IgG and any bound phage are captured by immunoprecipitation using protein A/G coated magnetic beads. The VSG-encoding inserts from bound phage are PCR amplified and sequenced. **(B)** Treemap of trypanosome subspecies and N-terminal domain type composition of VSGs included in the design of our phage display library. N-terminal domain types occur at similar frequencies regardless of species origin. **(C)** Histogram of the distribution of detected phage clones within the final library.

Sequencing of the phage display library revealed a normal distribution of inserts, though only 66.6% of tiles in the original library were detected (Fig. 1C). While PhIP-seq assays in the literature aim for at least 90% detection, these phage display libraries have also undergone less stringent clustering than ours and feature diverse sets of antigens belonging to many organisms and protein families. Ours, on the other hand, includes an inherently redundant set of antigens composed exclusively of VSGs, which are known to generate diversity through homologous recombination and often share similar peptide sequence with other variants. We used a network graphing approach^24,28^ to determine how well the 66.6% of the pool that remained after cloning and expansion represented the antigenic space covered by the original pool of 76,601 tiles. This analysis revealed that the loss of unique antigenic specificities was in fact minimal, and that 84.7% of the sequence in the original design could be found in its entirety within the cloned phages (Supp. Fig. S1A).

Our goal in designing the VSG phage library was to make it as diverse and representative as possible. However, antigens from parasites isolated from natural infections are underrepresented in the publicly available VSG databases used to design our library. Therefore, we sought to determine whether the library was sufficiently representative of the VSG repertoires present in human-infectious trypanosomes in the field. To do this, we assembled and identified VSG ORFs *de novo* from short-read whole genome sequencing data for 5 strains of *T. b. brucei*, 9 of *T. b. rhodesiense*, and 106 of *T. b. gambiense* type 1 and 6 of *T. b. gambiense* type 2 (all patients in the gHAT cohort were infected with *T. b. gambiense* type 1). Confirming the accuracy of the assembly, the majority of VSGs identified in the *T. brucei* Lister 427 laboratory strains had a top hit of > 90% identity within the existing genomic reference assembly for this strain (Supp. Fig S1C). We then compared assembled VSGs from all species to detectable phage insert sequences present in our library, which suggested that most of the VSGs in circulating strains have highly homologous counterparts within the final VSG phage display library (Supp. Fig. S1B). Based on these quality control analyses, we determined that the VSG phage display library was sufficiently representative of the *T. brucei* VSG repertoire, including that of human-infective *T. brucei* parasites in the field.

### Seroprevalent VSG peptides can be identified using PhIP-seq, though the anti-VSG IgG response is largely patient-specific

Using the VSG phage display library, we sought to broadly characterize anti-VSG antibody responses in a large collection of HAT patient serum samples and endemic control serum samples from the WHO HAT Specimen Biobank^29^. Samples were collected from *T. b. gambiense* patients in the Democratic Republic of Congo (n = 80), Guinea (n = 12), or Chad (n = 8). Negative controls for this gHAT cohort were individuals living in the endemic area that were both CATT and trypanolysis negative (DRC n = 81, Guinea n = 13, Chad n = 6). *T. b. rhodesiense* samples were collected from patients in Malawi (n = 58), Uganda (n = 11), and Tanzania (n = 1). Endemic negative controls were individuals living in the endemic area that were negative by both CATT and trypanolysis and had no parasites detected microscopically after concentration by capillary tube centrifugation (CTC) technique (Malawi n = 25, Uganda n = 0, Tanzania n = 25). Infected persons are classified as disease stage 2 based on presence of trypanosomes detected microscopically in CSF or elevated WBC counts (> 6 WBC/μL CSF). Those who are positive but with no evidence of CNS infection are designated as stage 1. In addition to HAT and endemic control serum samples, we also analyzed non-endemic control sera. Demographic information collected for study participants is presented in Table 1. In total we analyzed sera from 100 gHAT positive individuals, 100 gHAT negative endemic controls, 70 rHAT positive individuals, 50 rHAT negative endemic controls, and 36 non-endemic negative controls.

**Table 1).**
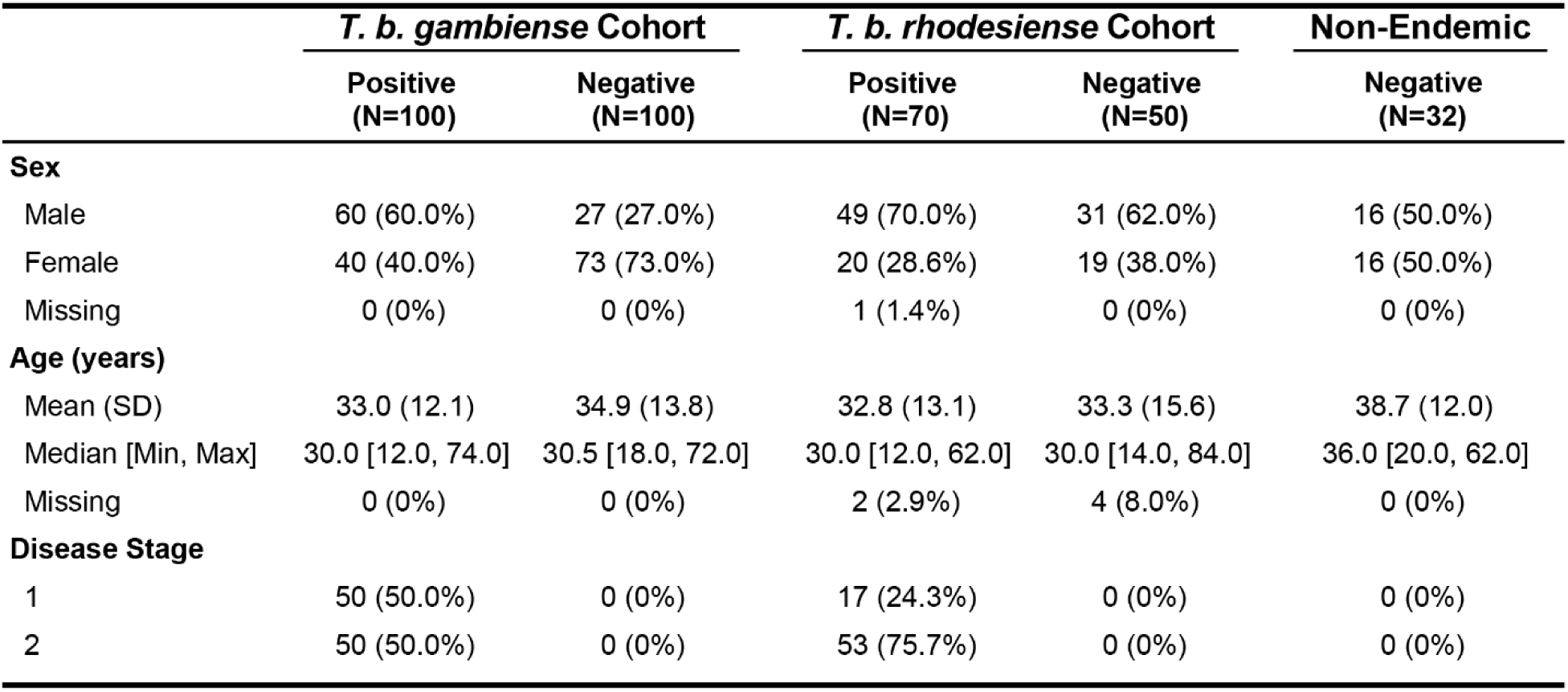
Study participant demographic data.

To investigate the breadth of the adaptive immune response against VSG peptides, we performed immunoprecipitation (IP) with protein A/G beads, which preferentially bind IgG. Peptide seropositivity was determined using an established approach where the signal in each sample is compared, using EdgeR, to a set of “mock” IPs which are run without serum. These mock IPs should reflect the baseline representation within the library as well as non-specific binding to the protein A/G beads. Consistent with a VSG-specific antibody response, we found that significantly more peptides enriched in stage 1 and 2 gHAT patients than in matched endemic negative controls (p = 1.5e-10 and p = 5.4e-11 respectively), with a higher number of peptides enriched in stage 2 than stage 1 patients. Conversely, only in stage 1 rHAT patients was the number of enriched peptides significantly higher than the corresponding endemic negative controls (p = 0.0071). Non-endemic control samples consistently generated the least signal (about 500 peptides per sample on average) (Fig. 2A).

**Fig. 2).**
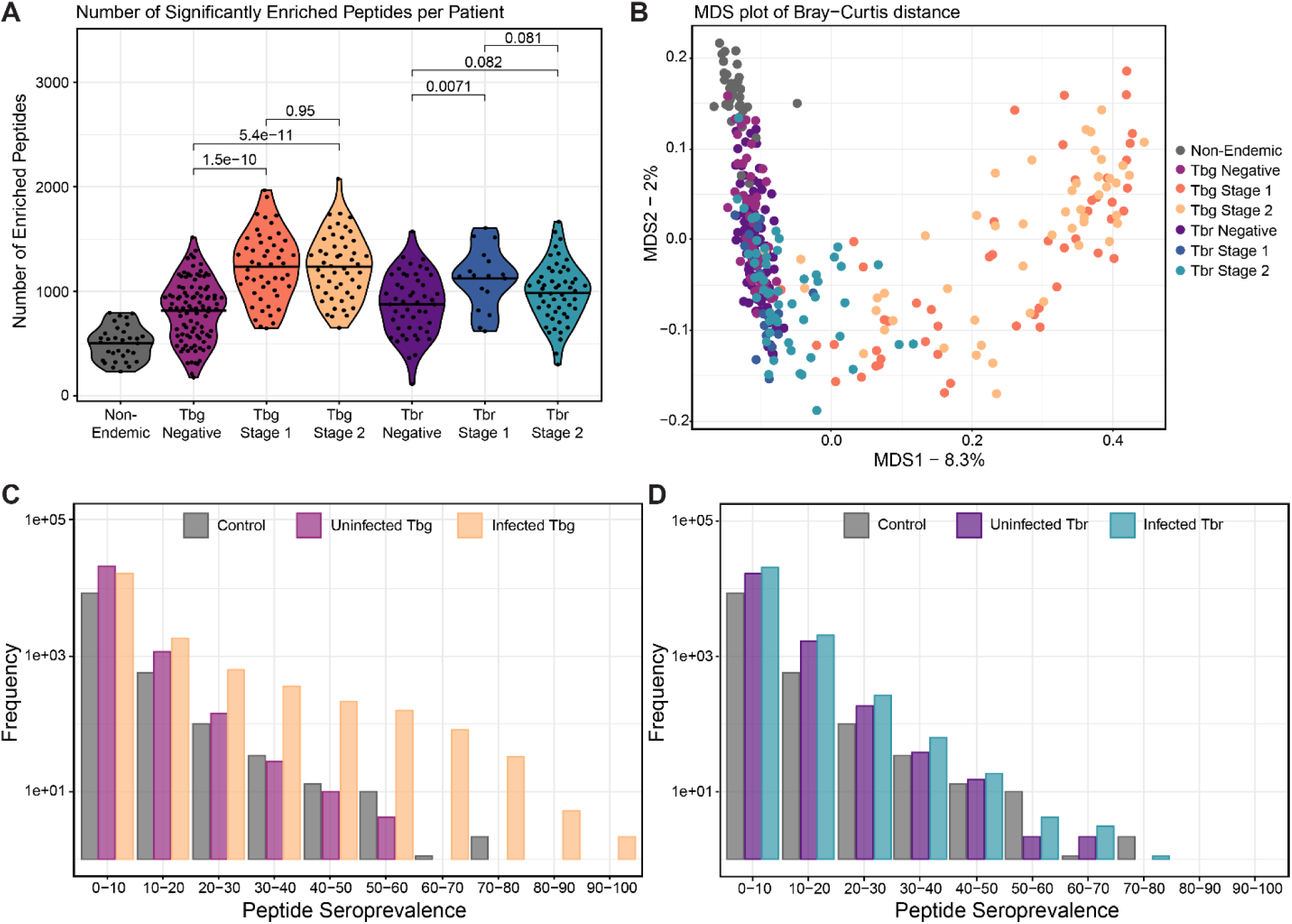
Profiling α-VSG IgG antibody epitopes in gHAT and rHAT cases. **(A)** Comparisons of the number of significantly enriched VSG peptides between patient groups. P-value statistics compare group means by Wilcoxon ranked sum test. All patient groups differ significantly from Non-Endemic Controls. **(B)** Multidimensional scaling analysis of the Bray-Curtis Dissimilarity calculated using Log_2_FC values, that is the amplitude of peptide enrichment per sample over the mock controls with no added serum. **(C-D)** Histograms showing counts of significantly enriched peptides binned by seroprevalence (bin size = 10%).

While anti-VSG antibody responses were largely similar within each patient group in terms of the number of unique antigens bound, counting peptides provides little information about the similarity of sets of VSG peptides bound by antibody across patients. To visualize the heterogeneity of anti-VSG IgG responses between individuals, we performed multidimensional scaling (MDS) analyses by calculating Bray-Curtis dissimilarity between patients. These visualizations suggest that the peptide reactivities in gHAT patients are more distinct from endemic controls than those of rHAT patients, though there was no clear clustering that distinguished between stage 1 and 2 gHAT (Fig. 2B). Furthermore, individual profiles of anti-VSG specificities do not appear to be meaningfully affected by sample collection sites or subjective measures of disease severity (Supp. Fig. S3, S4).

To further probe overlap in the anti-VSG specificities in each patient, we sought to identify seroprevalent peptides. These peptides are of particular interest since they indicate the presence of common antibody specificities among patients, either in response to the same antigen or to some conserved feature of the VSG. We found that antibody responses are mainly individual-specific: only 3,257 of the ∼30,000 total enriched peptides were shared between more than 10% of gHAT patients (Fig. 2C). However, several peptides proved to be highly conserved across patients, with some enriched in more than 80% of the infected cohort. Notably, we identified two peptides that are enriched in more than 90% of gHAT samples. In *T. b. rhodesiense* patients, on the other hand, a smaller proportion of peptides were shared between patients and the maximum seroprevalence was 78% (Fig. 2D). Seroprevalent peptides were detected in patients from all sites, suggesting they were not associated with certain geographical regions (Supp. Table S1). These did not share any primary peptide sequence similarity with either of the most commonly used diagnostic VSGs (LiTat1.3 and LiTat1.5) and thus represent novel antigens frequently bound by antibodies generated in response to *T. b. gambiense* infection that may be useful for diagnosis.

### PhIP seq reveals targets of anti-VSG immunity in gHAT and rHAT

Next, we sought to use the PhIP-seq enrichment data to understand how VSGs are targeted by the host immune system. We classified the NTD type for all sequences used in the design of VSG peptide inserts, extracted the coordinates of the NTD-CTD boundaries, and determined the origin of each NTD peptide tile. Our NTD identification pipeline was able to determine the type for nearly all (92.6%) detected library tiles of NTD origin. An analysis comparing the relative frequencies of peptide NTD types in seropositive peptides from infected cohorts to the frequency of NTD types in the phage library prior to enrichment showed that enriched peptides differed significantly from phages (χ^2^ p = 2.2e-16), with the gHAT cohort making the largest contribution to this difference (Fig. 3A). Most of the seropositive peptides were derived from the NTD in the gHAT cohort, while enriched peptides in the rHAT cohort largely resembled the composition of the phage library.

**Fig. 3).**
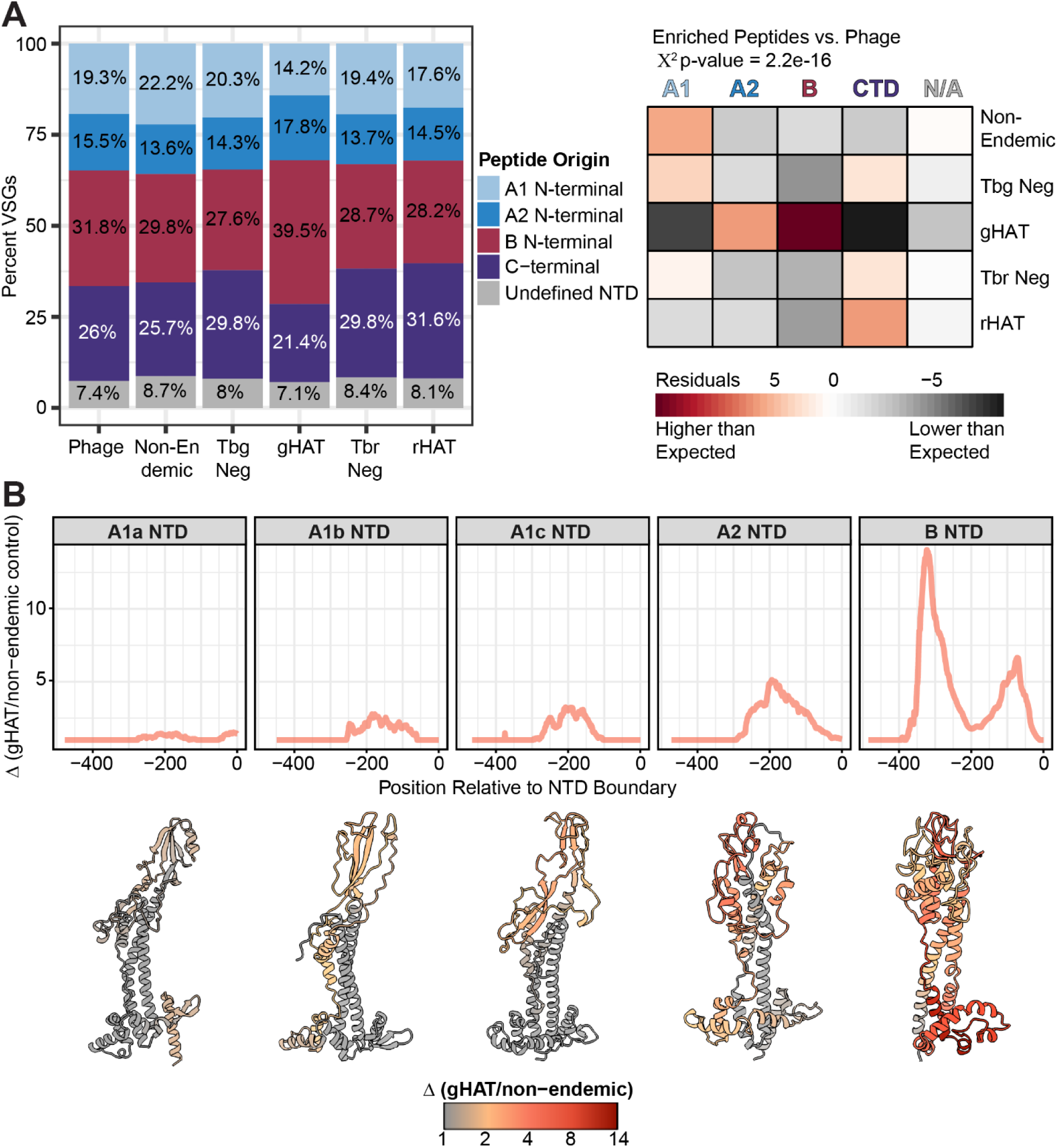
PhIP-seq reveals common targets of anti-VSG immunity. **(A)** Stacked bar charts quantifying the representation of VSG CTD tiles and NTD tiles of each type in the starting library and post-enrichment with *T. b. gambiense* or *T. b. rhodesiense* infected and negative control serum samples. Corresponding heatmap shows the residuals of χ^2^ statistical analysis of enriched groups vs. the expected frequency of peptides in the phage library. **(B)** Quantification of anti-VSG reactivity of antibodies in the serum of gHAT patients plotted along VSG NTDs of each type with 0 indicating the CTD/NTD boundary. Reactivity is calculated as delta (Δ), or the fold change in enrichment signal between the gHAT cohort and non-endemic controls which are assumed to be nonspecific. The representative models of NTD monomers shown below each plot facet are the most seroprevalent complete NTD of that type within the gHAT cohort. Models are colored on a gradient scale of gHAT signal over non-endemic controls.

Peptide identity itself provides little information regarding immunogenic regions of the VSG protein. The 90 residue tiles, while likely retaining some secondary structure, cover nearly a quarter of a complete VSG protein. B-cell epitopes, on the other hand, consist of 15 residues on average^30^. Because sets of peptides with very similar amino acid sequences frequently co-enrich together in multiple patients, it is possible to identify the most probable epitope containing sequences within the enriched tiles in a single patient. By defining the region of shared sequence among groups of related enriched peptides within an individual patient, we defined the most likely epitopes bound by each patient’s antibodies. Our primary focus was on the VSG epitopes targeted in gHAT patients, as the limited clustering and relatively low number of enriched peptides in rHAT suggests that much of the signal in these patients was non-specific. Once probable epitopes were defined for each patient, we mapped these sequences to a reference set of complete VSG protein sequences. The coordinates of mapped epitopes for each patient were summarized according to NTD type, allowing us to determine the most frequently targeted regions of each VSG. In gHAT, the highest enrichment was consistently seen within the top lobe of the VSG at the points most distal from the site of plasma membrane attachment, although foci of relatively high antibody binding are also observed on outer helices of the stalk as well as the bottom lobe of some NTD types (Fig. 3B). Despite the low specific signal in rHAT patients, NTD epitopes identified in this infection showed similar patterns (Supp. Fig. S5).

### Antibodies from gHAT patients preferentially bind peptides derived from type B VSGs

The most striking pattern that emerged from our enrichment data was a pronounced bias in enrichment towards peptides originating from type B NTDs in the gHAT patient cohort. The proportion of enriched type B peptides was significantly greater for gHAT disease stages 1 and 2 (p = 1e-11 and p = 5e-16 respectively) than their endemic uninfected controls, while all other sample groups bound to type B peptides at a frequency roughly equivalent to the distribution of type B peptides within the phage display library (Fig 4A).

**Fig. 4).**
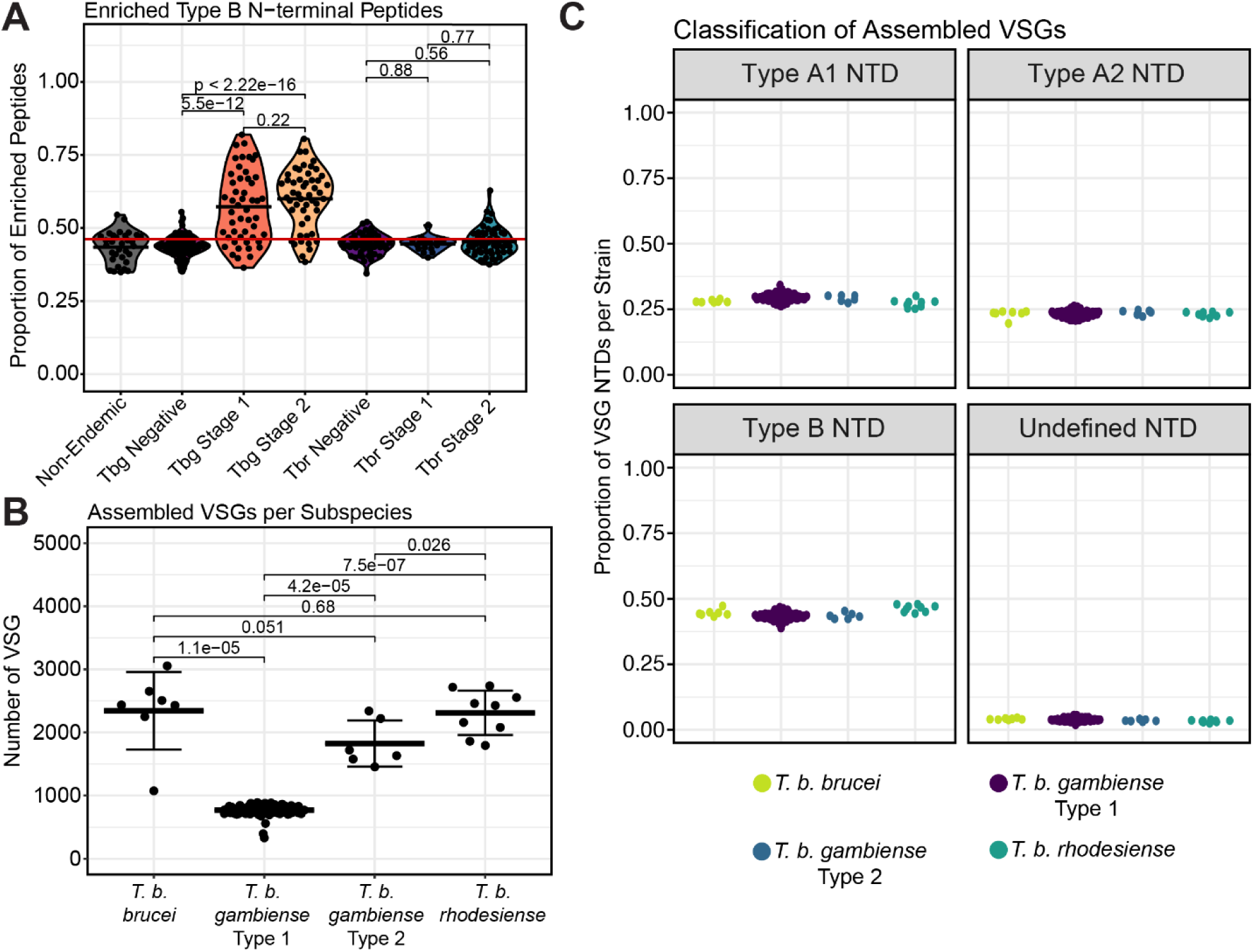
B-type VSGs are evenly represented across subspecies genomes. **(A)** The proportion of enriched type B NTD peptides per patient. Statistics compare group means by Wilcoxon ranked sum test. The red line denotes the proportion of NTD tiles in the starting library that are type B. **(B)** Whole genome short reads for *T. b. brucei* (n=7), *T. b. gambiense* type 1 (n=100), *T. b. gambiense* type 2 (n=6), and *T. b. rhodesiense* (n=9) field and laboratory strains were acquired from Data Dryad, ENA, and SRA databases. VSG repertoire size for each strain was estimated by assembling and identifying VSG genes using the VSG-seq methodology, calling with a minimum length of 300-AA and collapsing ORFs with 100% identity. **(C)** The proportions of VSG N-terminal domain types in the genomic repertoires of each strain. The group means of each subspecies do not differ significantly.

A bias towards expression of B type VSGs in *T. b. gambiense* could explain the predominance of type B VSG-derived peptides bound by gHAT patient sera. Such an expression bias could arise from the underlying composition of the *T. b. gambiense* VSG repertoire, which could contain more type B VSGs than other *T. brucei* subspecies, thus making these parasites more likely to elicit a predominantly anti-B VSG antibody response. Unfortunately, the VSG repertoire of *T. b. gambiense* is poorly characterized and it is unclear whether it contains a predominance of type B VSGs. To investigate the possibility that the predominance of type B-binding antibodies in gHAT patient sera was the result of differences in the underlying composition of the VSG repertoires of each *T. brucei* subspecies, we classified N-terminal domain types and compared the VSG sequences assembled from 126 whole genome sequencing datasets representing *T. b. brucei, T. b. rhodesiense,* and both types of *T. b. gambiense*.

The VSG repertoires of *T. b. brucei, T. b. rhodesiense,* and *T. b. gambiense* type 2 share many similar VSG sequences and cluster tightly together away from *T. b. gambiense* type 1 (Supp. Fig. S2A). Thousands of unique VSG ORFs assembled for strains of *T. b. brucei* (av. 2343.74), *T. b. rhodesiense* (av. 2309.89), and *T. b. gambiense* type 2 (av. 1824.33), while for *T. b. gambiense* type 1 the average number for type 1 was 769.59, confirming previous estimates that the *T. b. gambiense* type 1 VSG repertoire is smaller^31,32^ (Fig. 4B). We could identify both geographic and temporal clustering of VSGs from *T. b. gambiense* type 1, further supporting the accuracy of the assembly (Supp. Fig. S2B-C). Notably, the frequencies of assembled NTD types were consistent across different subspecies, with roughly 50% of VSGs being type B (Fig. 4C). This suggests that the observed bias, with antibody responses predominantly targeting type B peptides in gHAT patients, is unlikely to be the result of an expression bias driven by composition of the genomic VSG repertoire in *T. b. gambiense.* While a bias towards expression of B type VSGs might still exist in *T. b. gambiense* parasites despite a similar repertoire composition, an alternative explanation exists relating to the serological test used to screen individuals in this study. This test uses a parasite line expressing the VSG LiTat1.3, a type B VSG; thus, the CATT may preferentially detect infection in individuals with antibodies targeting B type VSGs.

### Reactivity against subsets of peptides can accurately distinguish infections and controls

If the CATT were biased towards detecting infection in individuals with anti-B VSG antibodies, it would be important to identify new diagnostic antigens for future serological tests. With this in mind, we sought to determine whether antibodies against seroprevalent peptides, particularly in combinations that include both A and B type VSGs, could serve as diagnostic markers of infection. We initially compared population-level differences between infected and endemic uninfected cohorts by significance testing to identify a long list of candidate diagnostic peptides. We then narrowed down this subset further using the recursive feature elimination method in the caret R package^33^ to remove peptides with the weakest association with infection. Next, we fitted models for a variety of machine learning methodologies using peptide enrichment values (Log_2_FC of samples vs. mocks) as predictors. We found the optimal models for each method by repeated k-fold cross-validation and selected those with the highest test set prediction accuracy in classifying samples as infected or uninfected. Performance was comparable across all machine learning methods used for each classification (Fig. 5A-B). The models demonstrated remarkable accuracy in distinguishing the serum of rHAT patients from their healthy endemic counterparts, with area under the curve (AUC) values ranging from 0.79 to 0.87 (Fig. 5A). Classification accuracy for distinguishing gHAT patients from endemic controls was even higher with AUC values ranging from 0.95 to 0.99 (Fig. 5B). Top models for predicting infection status had F1 scores >0.76 for rHAT and >0.97 for gHAT. This analysis demonstrates that measuring antibody reactivity against VSG peptides can accurately distinguish between infected and uninfected serum samples for gHAT as well as rHAT.

**Fig. 5).**
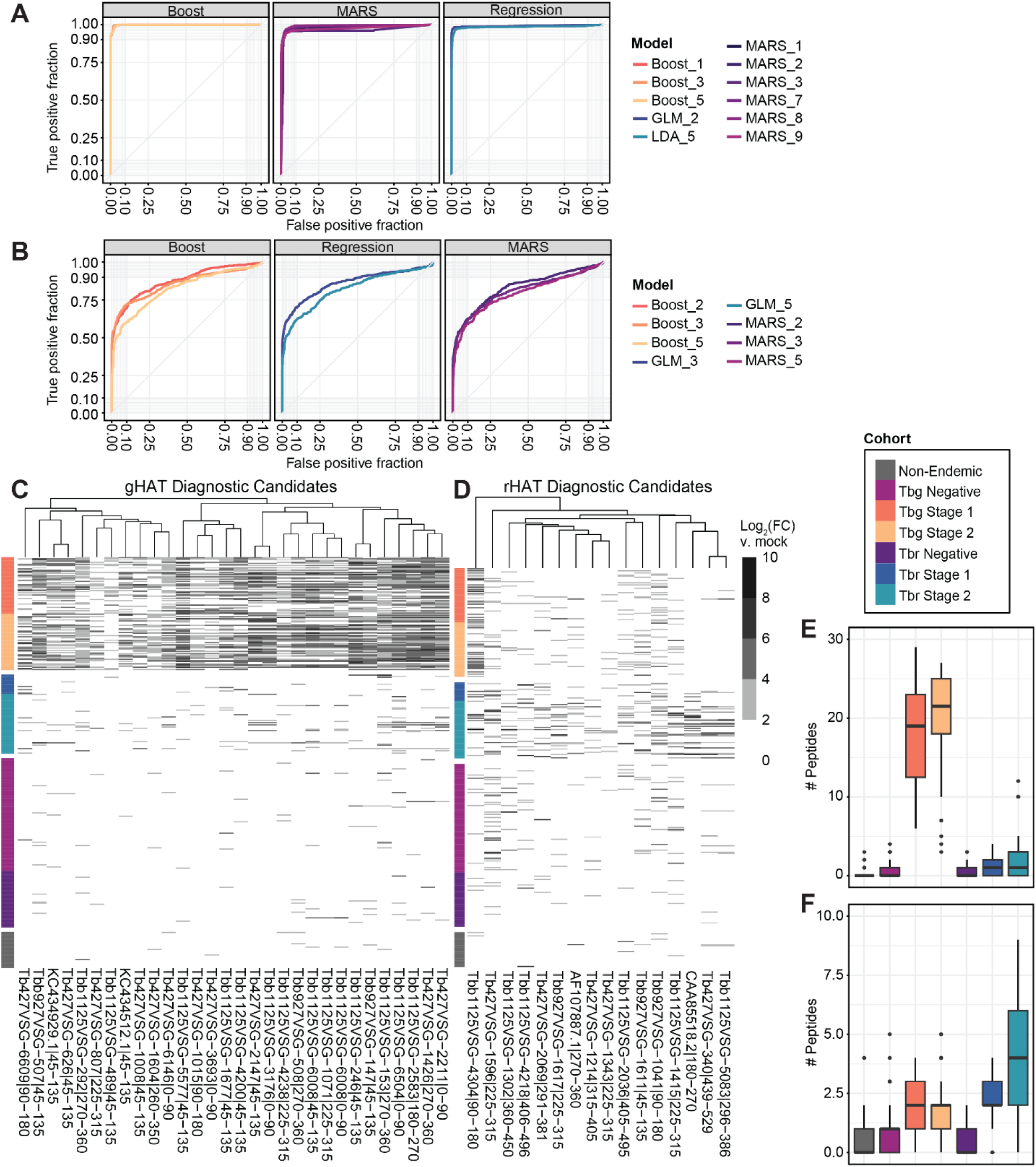
Panels of potential gHAT and rHAT diagnostic peptides. ROC curves demonstrate the efficacy of the best performing cross-validated models for classifying **(A)** *T. b. gambiense* and **(B)** *T. b. rhodesiense* infected individuals within the testing dataset (20% of samples). **(C-D)** Hierarchically clustered heatmaps of IgG reactivity to discriminatory peptides identified by the statistical machine learning models shown in (A) and (B). The identities of peptide tiles are shown as column names. The proxy for antibody reactivity is Log_2_(FC) enrichment vs. mock beads-only immunoprecipitation reactions which had no added serum. **(E-F)** Quantification of the number of gHAT or rHAT peptides respectively that could be robustly detected with PhIP-seq (Log_2_(FC) ≥ 2) per sample.

A new serological diagnostic that could be practically applied in endemic areas would ideally detect antibodies against one or a few peptide antigens, fewer than the long list of peptides produced by the models. To identify the best candidate peptides to potentially develop as diagnostic targets, we computed the relative variable importance of each peptide in top performing models (Supp. Fig. S6A, S7A). The reactivity profiles of peptides with high importance in the most accurate models were visualized (Fig. 5C-D). The candidate diagnostic peptides demonstrated high specificity, with minimal or sporadic reactivity detected in the endemic uninfected cohorts. On average, at least 2 of the total 16 rHAT predictive peptides are robustly detected by the PhIP-seq assay with fold change ≥ 4 in the serum of *T. b. rhodesiense* infected individuals (Fig. 5E). Notably, closer inspection of these top diagnostic candidates revealed that two of the peptides, although annotated as VSGs in public databases, did not in fact originate from VSGs. These peptides belong to known virulence factors ISG65 (AF107887.1) and SRA (CAA85518.2), both of which are invariant proteins expressed on the surface of *T. b. rhodesiense*. This suggests the invariant proteins, rather than VSGs, may serve as even more sensitive markers of rHAT infection. For gHAT, on the other hand, we identified 30 possible diagnostic peptides, all of which appear to derive from VSG and are highly enriched in the serum of *T. b. gambiense* infected patients (Fig. 5F).

Since VSGs have the potential to diversify, compromising the effectiveness of any VSG-based serological test, we also assessed the likelihood of diagnostic candidates to be maintained in the genomes of circulating strains. By searching for similar sequences within VSG repertoires assembled from field isolates, we found that many of the gHAT and rHAT candidate peptides could be found in genomes of the same subspecies (Supp. Fig. S6B, S7B). Overall, our analysis shows PhIP-seq is a viable method to screen antibody reactivities against thousands of peptide antigens as an unbiased approach for identifying potential targets for serological diagnostics. Moreover, these data show the potential promise of using combinations of peptide antigens, whether VSG-derived or otherwise, for the serological diagnosis of HAT.

## Discussion

Although antigenic variation of the VSG is *T. brucei*’s primary mechanism of maintaining a chronic infection, little is known about the VSG-antibody interface, with human antibody responses to VSG remaining virtually unstudied. What is known about these interactions has largely been generalized from the results of low-throughput experiments focused on handfuls of VSGs, or from mouse models and the animal-infective *T. b. brucei*. Here, we overcome previous technical hurdles and characterize, for the first time, the antibody response to VSG in hundreds of HAT patients. Using PhIP-seq, we evaluated antibody binding to tens of thousands of VSG peptides, revealing important differences between *T. b. rhodesiense* and *T. b. gambiense* infection. Our analysis allowed us to identify the parts of the VSG protein targeted by antibody and the specific epitopes bound. Despite finding that anti-VSG repertoires were largely patient-specific, we were also able to identify candidate antigens for serological HAT diagnostics, which could play an important role in HAT elimination.

An overarching theme in our analysis was the stark difference between the antibody responses against *T. b. gambiense* and *T. b. rhodesiense*. While clear anti-VSG responses, distinct from endemic controls, could be detected in the gHAT cohort, rHAT patients had relatively few seroprevalent peptides and clustered with endemic controls in MDS visualizations. We hypothesize that this difference is related to the unique characteristics of each infection. *T. b. gambiense* has a smaller VSG repertoire than *T. b. rhodesiense* and causes a chronic infection characterized by low parasitemia, while *T. b. rhodesiense,* with a larger VSG repertoire, causes an acute infection characterized by high parasitemia^31,34^. This combination of a smaller antigenic repertoire and a smaller antigen load due to low parasitemia may allow for the development of a memory response to VSG and high-affinity antibodies, with specificities accumulating over the course of infection. On the other hand, the acute nature of *T. b. rhodesiense* infection may lead to a predominance of lower affinity IgM antibodies, which cannot be detected by our assay, which uses IgG-bidning Protein A/G beads. This is supported by the fact that IgG antibodies against invariant proteins, like ISG65 and SRA, were consistently detected in rHAT patients. These antigens, which the host is continuously exposed to, may allow for the elicitation of the high affinity IgG responses that are preferentially detected in our assay.

It is also possible that the difference between gHAT and rHAT antibodies we observe could be related to the VSG antigens recognized in each infection. While our use of long peptides in the library maintained much secondary structure, the T7 bacteriophage display vector is unable to reproduce post-translationally modified or conformational epitopes. If *T. b. rhodesiense* N-terminal epitopes are predominantly conformational or post-translationally modified, these would be missed in our analysis, resulting in low specific signal. It will be critical to understand what proportion of the anti-VSG response is against post-translationally modified epitopes, as sugars on VSGs have been found to greatly affect the antigenic character of the VSG^35,15^. Alternatively, the low specific signal in rHAT patients could be the result of B cell memory loss, which has previously been observed in mouse models of *T. b. brucei* infection^36,37^, but not in *T. b. gambiense* patients, potentially reflecting the distinct antibody response associated with each infection^38^.

An alternative explanation for the overlap in specificities between rHAT patients and endemic controls could be exposure to the animal parasite *T. b. brucei* in endemic controls. Though this parasite is unable to establish infection in humans, patients in endemic regions are likely exposed to the VSGs of *T. b. brucei* routinely through bites of *T. brucei*’s tsetse fly vector. Evidence, including the repertoire analysis we report in this study, suggests that *T. b. rhodesiense* is essentially a variant of *T. b. brucei,* differing only by the presence of a single gene^39^. Therefore, antibody repertoires after exposure to *T. b. brucei* may be similar to those generated by infection with *T. b. rhodesiense*. These similarities between *T. b. brucei* and *T*. *b. rhodesiense* have perhaps contributed to difficulties in identifying antigenic, molecular, or serological targets for diagnosing rHAT^40–42^. Regardless of the explanation, it is clear that the antibody responses in gHAT and rHAT are distinct.

Using the clear and specific signal associated with gHAT infection, we were able to identify the VSG epitopes commonly bound by antibody during infection. While previous analyses of antibody binding to VSG were largely low-throughput, our analysis allowed us to evaluate these specificities in large numbers of patients for large numbers of VSGs. Our findings are consistent with previous studies of live-parasite-binding monoclonal antibodies and nanobodies which have suggested that antigenic surfaces can extend along the length of the VSG NTD^13,14^. Our high-resolution analysis confirms that these previous observations, made primarily in animal models, hold true in human infection as well. In gHAT infection, the top lobe of the VSG NTD is the predominant target of anti-VSG antibodies, although the entire VSG NTD can be immunogenic.

Our analysis of VSG epitopes revealed another striking, and unexpected, finding: the majority of anti-VSG NTD peptides in *T. b. gambiense* patient sera were derived from B type VSGs. This observation echoes our previous finding that type B VSGs are disproportionately expressed during *T. b. gambiense* infections, but not during *T. b. rhodesiense* infections^27^. Analysis of the VSG repertoire of 122 *T. brucei* field isolates suggests that this is not because *T. b. gambiense* carries more B type VSGs within its genome than the other subspecies. Indeed, all the parasite strains we analyzed had a near identical proportion of B type VSGs within their repertoires. While it remains possible that *T. b. gambiense* parasites preferentially express type B VSGs despite roughly equivalent numbers of each VSG subtype within the genome, this seems unlikely given the small genome and VSG repertoire of this subspecies. Why would the parasite with narrow host range retain a subset of VSGs in its minimal VSG repertoire that are infrequently used?

An alternative explanation seems more likely: in this study, as well as our previous analysis of VSG expression in *T. b. gambiense* patients, all infected patients were CATT-positive. The CATT uses parasites expressing the LiTat1.3 VSG, a type B VSG. It is therefore possible that that this test, which relies on a type B VSG for diagnosis, preferentially detects infection in patients infected with parasites expressing type B VSGs, who may be more likely to carry antibodies that cross-react with LiTat1.3. If this is the case, then gHAT may be underdiagnosed in populations where circulating parasites express different VSGs. It will be essential to evaluate both VSG expression and antibody responses in patients screened and diagnosed with tests that use different combinations of antigens, such as the newer RDTs for gHAT that include the type A2 VSG LiTat1.5 and ISG65 in addition to LiTat1.3 ^43^, to determine whether a bias in the CATT exists.

The possibility of screening tests introducing bias is important to consider as endemic nations work towards the elimination of HAT, as it could affect the ability to detect subsets of infected people. If such a bias exists, then it will be critical to identify unbiased diagnostic targets. While our sample set is not ideal for identifying unbiased diagnostic targets, because all patients were screened using the CATT, our analysis demonstrates the utility of PhIP-seq for identifying new HAT diagnostic targets. Using this high-resolution tool, we identified several candidate peptides associated with gHAT that bear no apparent peptide sequence similarity to either LiTat1.3 or LiTat1.5 and that represent both A and B type VSGs. These targets will require further validation with serum from patients who were screened using alternatives to the CATT.

Surveillance of rHAT would also greatly benefit from a serological diagnostic. Widespread adoption of rapid diagnostic tests for malaria has exacerbated rHAT underdiagnosis since there is now decreased infrastructure available for microscopic blood smear analysis^18^. For the first time, we have identified a small panel of peptides that elicit strong antibody responses specifically predictive of *T. b. rhodesiense* infection. Notably, two of the top-contributing peptides for classification of rHAT patients were not derived from VSG, but instead were invariant proteins expressed on the surface of bloodstream form parasites. Contrary to previous assumptions, our results suggest that a serological screen for rHAT is possible. However, they also indicate that the ideal diagnostic target for rHAT is likely not a VSG at all. Whole proteome PhIP-seq screens could prove extremely useful for identifying new serological targets for rHAT.

Altogether, our data provide a comprehensive picture of the antibody specificities that define human *T. brucei* infection. By defining the VSG epitopes bound by host antibody, we confirm the long-held assumption that the disordered regions at the top lobe of the VSG NTD are indeed a primary interface of antibody interactions. In addition to uncovering fundamental features of the antibody response to VSG, this study identified seroprevalent peptides for both rHAT and gHAT that could serve as the basis of new serological tests, which may be essential for gHAT elimination, if the CATT is biased as our data suggests. More broadly, this study underscores the utility of PhIP-seq as an approach to understand antibody-antigen interactions in the context of infectious disease. This high-resolution approach, where thousands of antibody specificities can be screened in hundreds of patients, is poised to play an important role in understanding how pathogens manage to continuously evade host antibody.

## Supplemental Figures

**Table S1).** Geographical Distribution of Seroprevalent (≥ 70%) Peptides in gHAT Cohort.

| Peptide | Guinea | DRC | Chad |
| --- | --- | --- | --- |
| KC434873.1\|225-315 | 9 | 62 | 4 |
| KC434929.1\|45-135 | 8 | 60 | 5 |
| Tb08.27P2.630\|74-163 | 9 | 55 | 7 |
| Tb09.v4.0189\|180-270 | 9 | 62 | 1 |
| Tb427VSG-1178\|45-135 | 8 | 59 | 6 |
| Tb427VSG-1294\|225-315 | 8 | 64 | 4 |
| Tb427VSG-1320\|45-135 | 9 | 59 | 6 |
| Tb427VSG-1426\|270-360 | 9 | 76 | 7 |
| Tb427VSG-1451\|180-270 | 9 | 61 | 5 |
| Tb427VSG-1697\|45-135 | 11 | 55 | 6 |
| Tb427VSG-1710\|45-135 | 7 | 57 | 6 |
| Tb427VSG-2147\|45-135 | 8 | 64 | 4 |
| Tb427VSG-2211\|0-90 | 10 | 78 | 7 |
| Tb427VSG-2371\|226-315 | 6 | 63 | 4 |
| Tb427VSG-3213\|45-135 | 9 | 64 | 6 |
| Tb427VSG-538\|45-135 | 10 | 54 | 6 |
| Tb927.3.410\|46-135 | 9 | 67 | 6 |
| Tbb1125VSG-1029\|45-135 | 6 | 70 | 1 |
| Tbb1125VSG-1071\|225-315 | 11 | 58 | 5 |
| Tbb1125VSG-153\|225-315 | 9 | 63 | 4 |
| Tbb1125VSG-1689\|90-180 | 7 | 57 | 6 |
| Tbb1125VSG-246\|45-135 | 9 | 64 | 5 |
| Tbb1125VSG-2583\|180-270 | 7 | 68 | 6 |
| Tbb1125VSG-292\|270-360 | 6 | 64 | 4 |
| Tbb1125VSG-3176\|0-90 | 8 | 62 | 2 |
| Tbb1125VSG-4060\|45-135 | 7 | 74 | 1 |
| Tbb1125VSG-4150\|135-225 | 9 | 58 | 3 |
| Tbb1125VSG-454\|90-180 | 7 | 60 | 5 |
| Tbb1125VSG-503\|45-135 | 4 | 62 | 6 |
| Tbb1125VSG-5509\|45-135 | 7 | 65 | 3 |
| Tbb1125VSG-5577\|45-135 | 8 | 59 | 7 |
| Tbb1125VSG-6008\|45-135 | 12 | 61 | 6 |
| Tbb1125VSG-6175\|0-90 | 6 | 61 | 3 |
| Tbb1125VSG-6504\|0-90 | 8 | 72 | 6 |
| Tbb927VSG-147\|45-135 | 9 | 62 | 5 |
| Tbb927VSG-2130\|159-249 | 8 | 58 | 4 |
| Tbb927VSG-508\|270-360 | 11 | 62 | 4 |
| TevSTIB805.11_03.470\|225-315 | 5 | 64 | 2 |
| TevSTIB805.7.7120\|378-468 | 8 | 66 | 6 |
| **Total Patients:** | **12** | **80** | **8** |

**Fig. S1).**
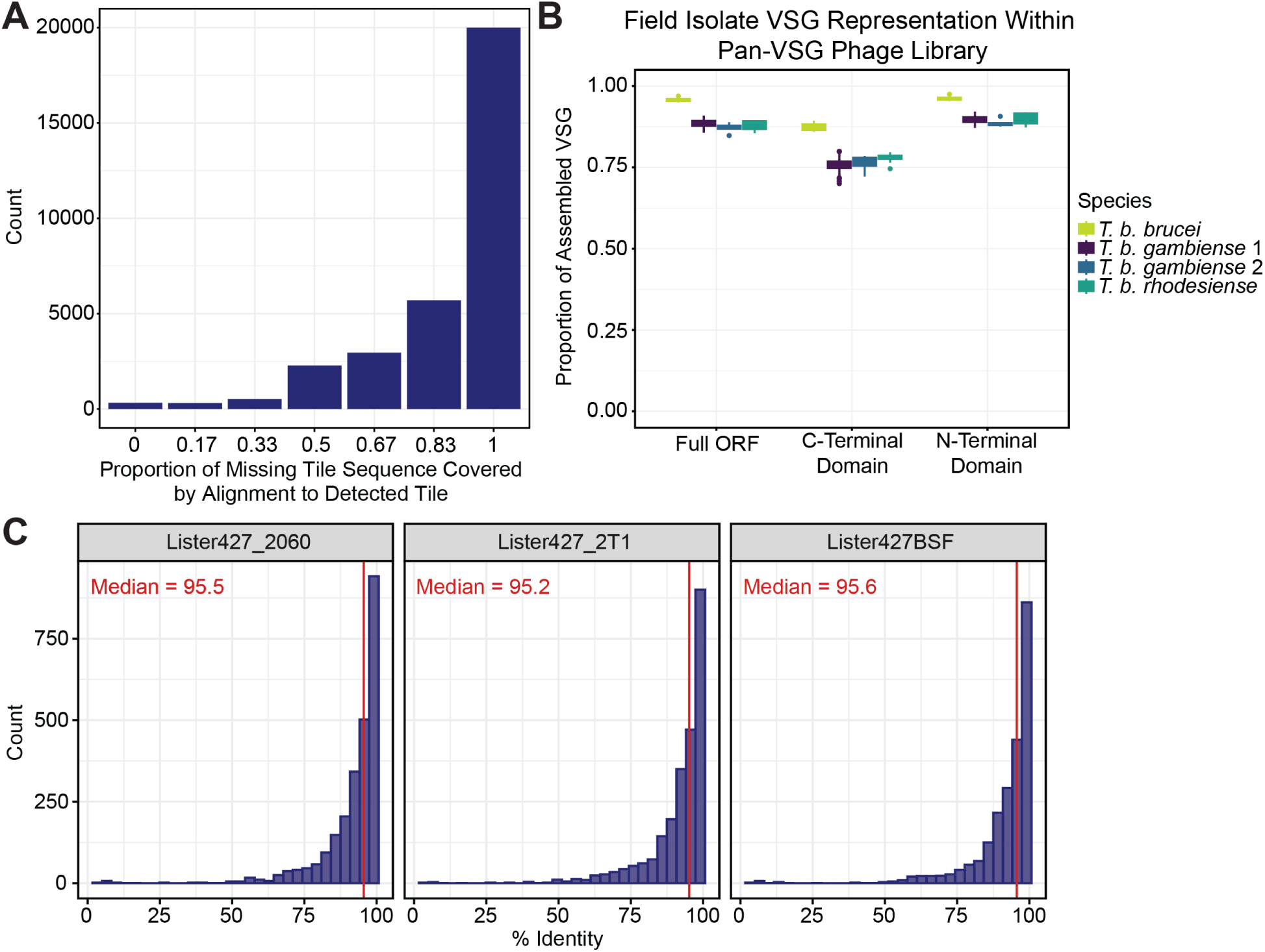
Phage Library approximates VSG repertoires of uncharacterized *T. brucei* isolates. **(A)** Histogram showing the counts of undetected peptide tiles with homologous sequence present in represented library tiles. 28,272 of 76,601 total peptide tiles were undetected. Of the missing tiles, 11,751 contained unique sequence that was not represented by library tiles, however this rarely constituted the entire tile. **(B)** Quantification of the proportion of assembled VSG per field isolate strain where the majority (≥80%) of the whole ORF, NTD or CTD sequence can be found within Pan-VSG library tiles. **(C)** Histograms showing the distribution of alignments between assembled VSGs from three Lister427 whole genome datasets and the complete Lister427 known VSG gene reference.

**Fig. S2).**
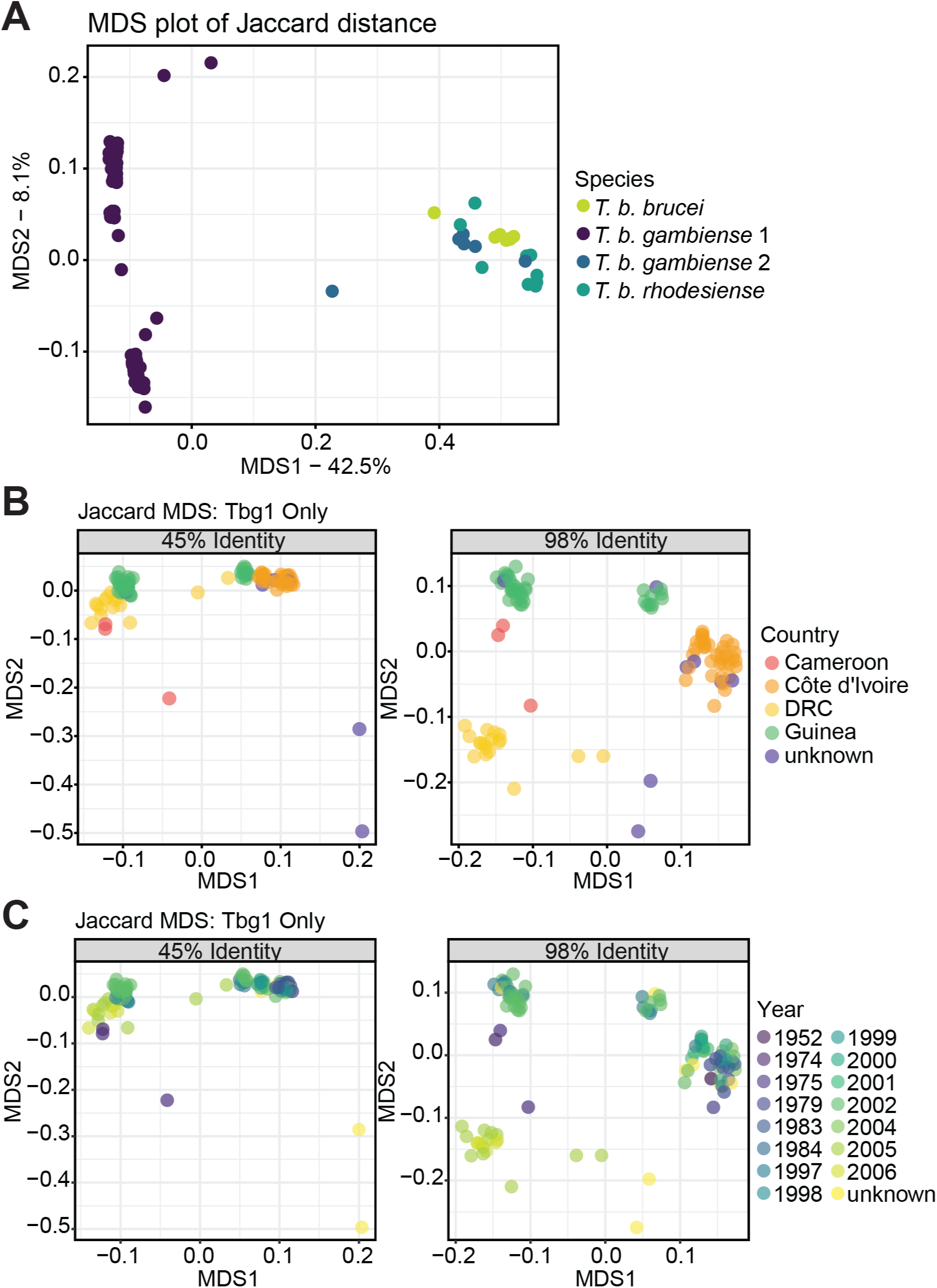
*T. b. gambiense* VSG repertoires are distinct from other *T. brucei* subspecies. (**A**) Multidimensional scaling analysis (MDS) of the Jaccard Distance calculated based on cluster membership. Assembled VSG from all strains were clustered at a global nucleotide sequence identity of 80% using cd-hit-est. **(B-C)** MDS analysis of the Jaccard Distance calculated based on peptide sequence clustering by cd-hit at global identity thresholds of 45% and 98%.

**Fig. S3).**
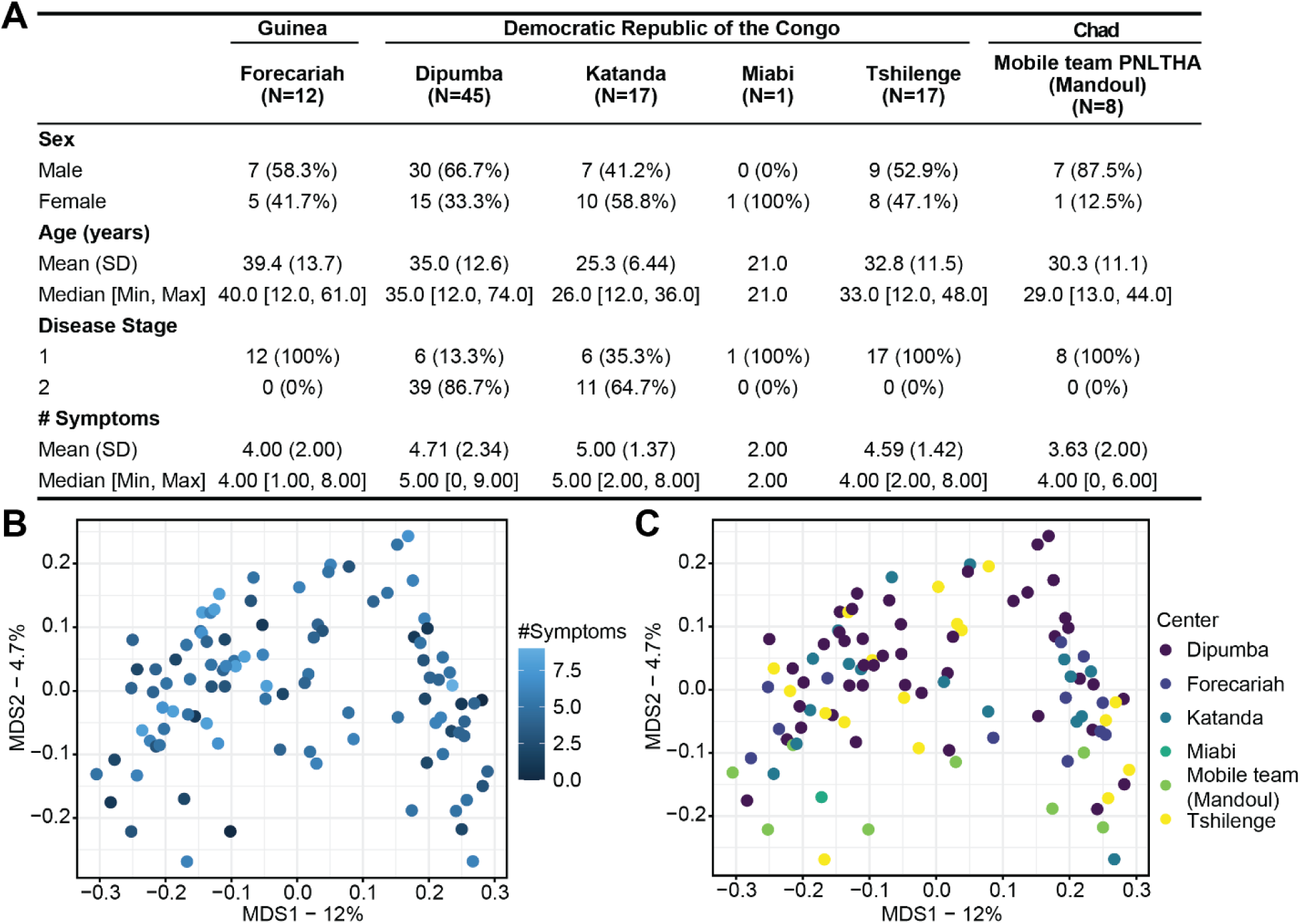
Associations between symptom severity and anti-VSG response in gHAT. **(A)** Demographic data for the T. b. gambiense infected cohort outlining relevant metadata for subjects at each sample collection site. Limited clinical data was collected for infected persons (including fever, faintness, weight loss, swollen lymph nodes, face edema, loss of appetite, and neurological signs) which was summarized as the number of symptoms reported. Multidimensional scaling analysis of the Bray-Curtis Dissimilarity calculated based on enrichment (Log_2_(FC)) of peptides in the serum of T. b. gambiense infected individuals colored by the number of recorded symptoms **(B)** and the collection site **(C)** does not reveal any strong drivers of clustering between samples.

**Fig. S4).**
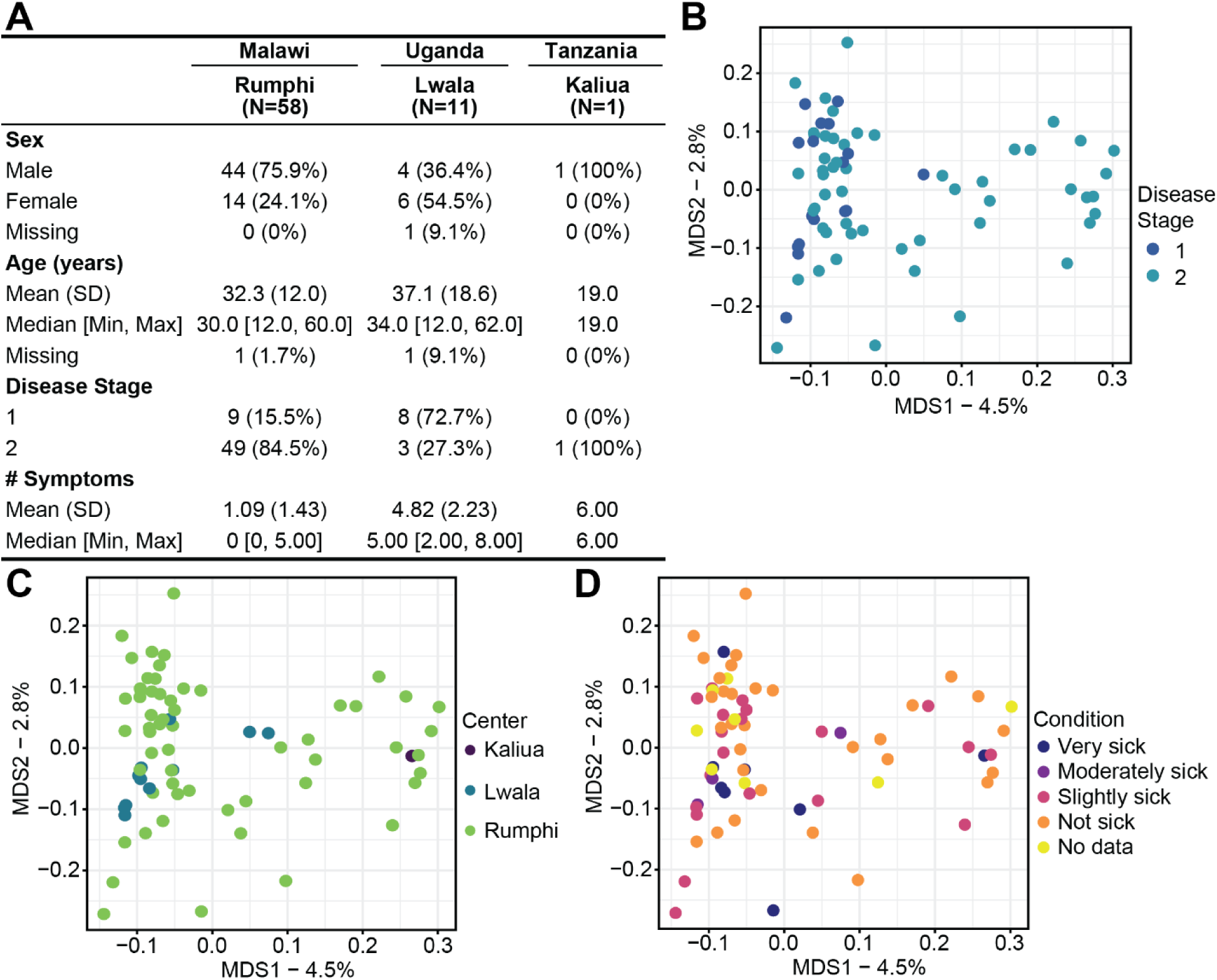
Associations between symptom severity and anti-VSG response in rHAT. **(A)** Demographic data for the T. b. rhodesiense infected cohort outlining relevant metadata for subjects at each sample collection site. Most clinical symptoms which were recorded at other sites were not included in the Malawi Rumphi sample collection, however, qualitative assessment of physical condition was reported. **(B-D)** Multidimensional scaling analysis of the Bray-Curtis Dissimilarity calculated based on enrichment (Log_2_(FC)) of peptides in the serum of T. b. rhodesiense infected individuals.

**Fig. S5).**
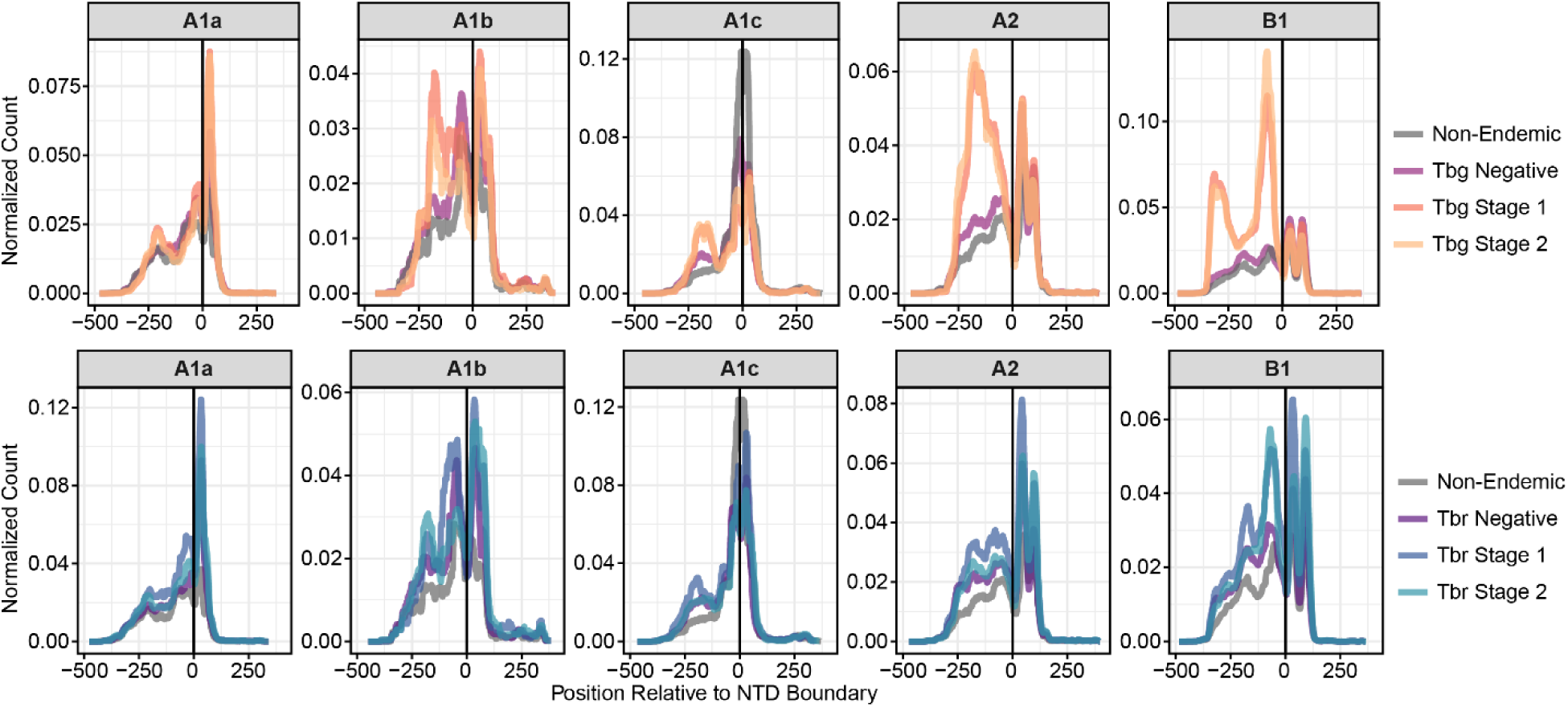
Immunogenic sites along the VSG. Enriched motifs mapped to the complete repertoire of full-length VSG sequences used to create the VSG phage library. Histograms for the gHAT and rHAT cohorts showing the abundance of enriched sequences mapped to VSGs of each NTD type, normalized by cohort size and the total number of VSGs of that type. The x-axis shows the residue position along each VSG protein relative to the NTD/CTD boundary which is at x = 0. A positive x coordinate falls within the NTD and positive values are within the CTD.

**Fig. S6).**
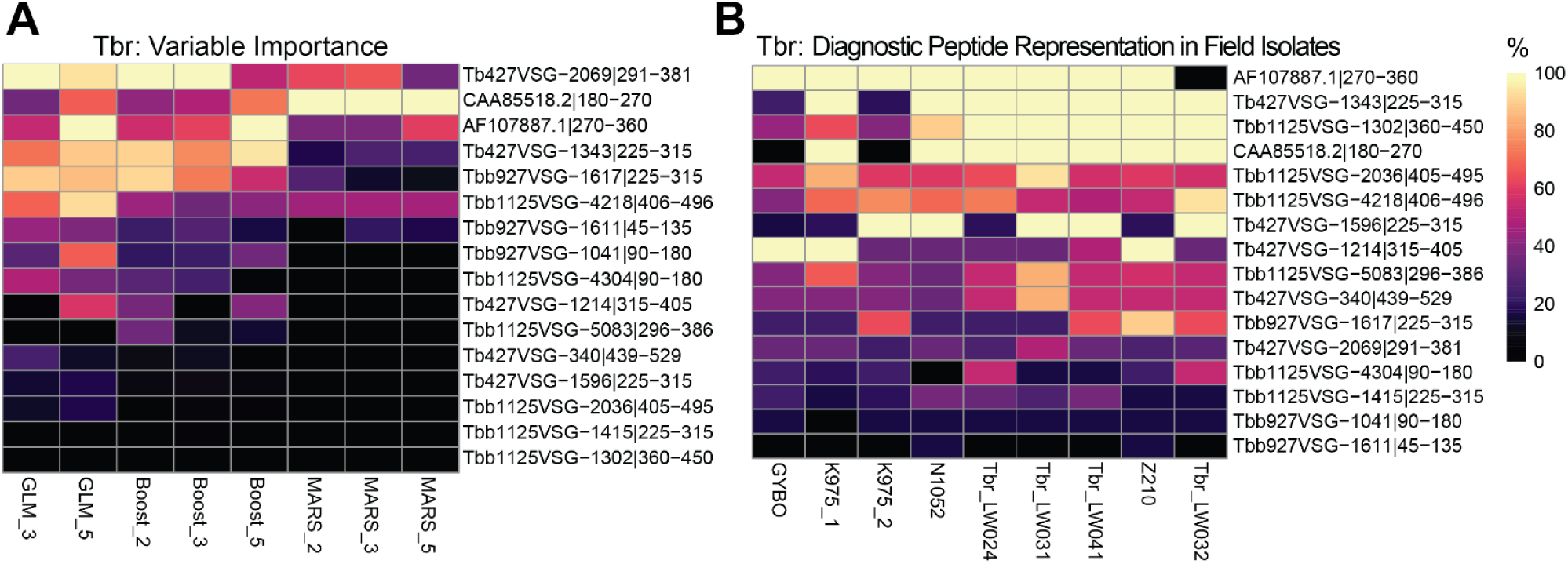
Ranking rHAT diagnostic peptide candidates. Heatmaps used for the ranking of candidate peptides for the diagnosis of *T. b. rhodesiense* infection ordered by cumulative row value. Peptide identity is labeled on the y-axis. Values are normalized and displayed as either **(A)** the percentage of variable weight attributed to a peptide by each model as indicated on x-axis, or **(B)** percentage of the peptide tile sequence that could be found within VSG proteins assembled from the field isolate genomes specified on x-axis.

**Fig. S7).**
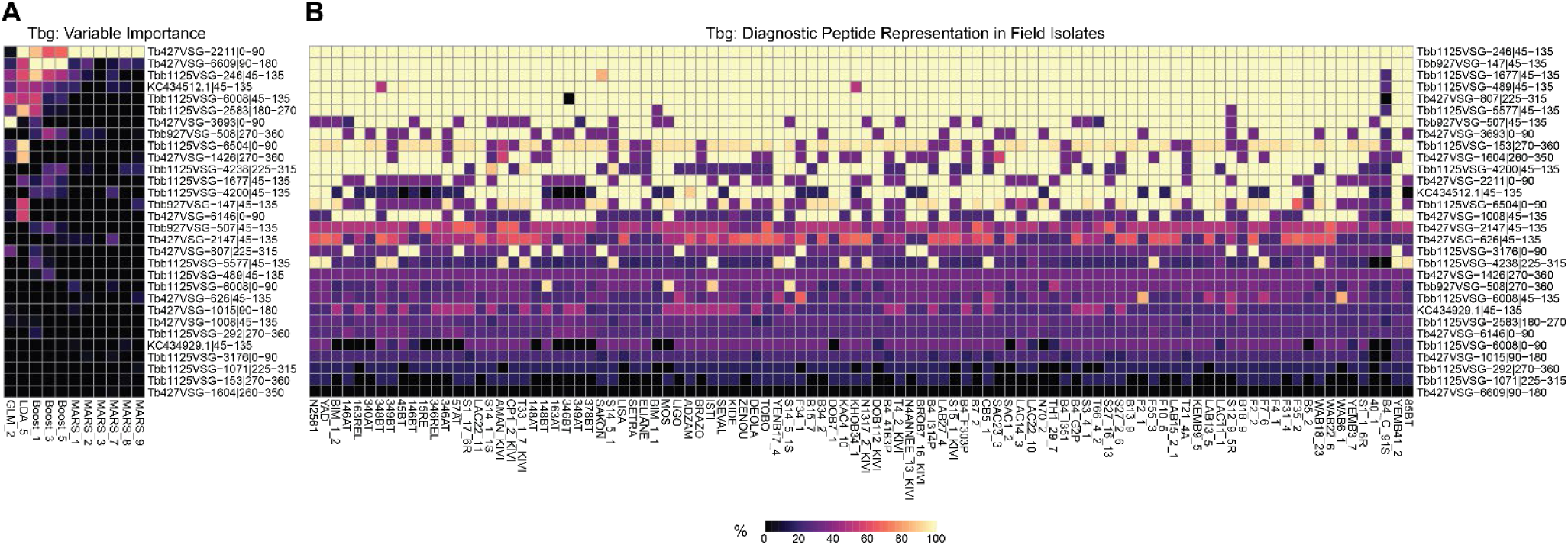
Ranking gHAT diagnostic peptide candidates. Heatmaps used for the ranking of candidate peptides for the diagnosis of *T. b. gambiense* infection ordered by cumulative row value. Peptide identity is labeled on the y-axis. Values are normalized and displayed as either **(A)** the percentage of variable weight attributed to a peptide by each model as indicated on x-axis, or **(B)** percentage of the peptide tile sequence that could be found within VSG proteins assembled from the field isolate genomes specified on x-axis.

## Methods

### Phage Display immunoprecipitation and Sequencing (PhIP-Seq)

**(i)** **Library Design and Cloning.** The VSG phage display library is intended to cover all variant surface glycoprotein (VSG) antigens expressed by the human-infectious *T. b. gambiense* and *T. b. rhodesiense* as well as the closely related non-human-infectious subspecies *T. b. brucei*, *T. evansi*, and *T. equiperdum*. Protein sequences were obtained from the curated VSG database at https://tryps.rockefeller.edu/Sequences.html as well as from Genbank and TriTrypDB by searching for entries annotated as VSG. Proteins were broken into large 90-amino acid (aa) peptide sequences with 45aa overlap between adjacent tiles to include as much secondary structure as possible. VSG pseudogenes were split up along their stop codons and tiled in between if the sequence was at least 90-AA long, while shorter fragments were discarded. The median length of discarded pseudogene fragments was 9aa. Then we collapsed all 100% identical tiles using the cd-hit algorithm (–c 1.0 –G 0 –A 90 –d 0) so only one representative was included in the final library design. By only clustering tiles at 100% identity, the library retains highly similar VSG sequences and represents the fullest diversity. We reverse-translated peptide tiles into DNA sequences codon optimized for expression in *Escherichia coli*, while synonymously recoding any EcoRI and HindIII restriction enzyme sites to facilitate cloning. Then we added to the 5’ end GTTAGCCGGAATTCC and to the 3’ AAGCTTGGGTGCTAC as adapters to add appropriate restriction sites for cloning into the T7 phage display vector, resulting in 300-nucleotide sequences which were synthesized by Twist Bioscience. Phage cloning was performed according to manufacturer guidelines using the T7Select 10-3 Cloning Kit (Sigma-Aldrich 70550-M).
**(ii)** **Phage Immunoprecipitation.** We screened serum IgG specificities against the Pan-VSG library following the previously published protocol^22^. In summary, we added 0.2 mL of a 100-fold dilution of serum sample to 1 mL of phage library solution containing 7.66 x 10^9^ pfu per reaction. The serum and phage were rotated overnight at 4 °C (∼16 h) to allow complex formation between phage and serum antibodies, then we added 20 μL each of protein A and protein G-coated magnetic beads (Invitrogen 100002D, 100004D) before rotating an additional 4 h at 4 °C. We washed the beads three times in a 0.3% NP-40 solution and then resuspended in a Herculase II (Agilent 600679) PCR master mix to amplify VSG inserts from bound phage genomes. This reaction underwent 18 cycles of PCR and 2 μL of product was added to a second PCR reaction with 22 cycles of amplification to add sample-specific barcodes and Illumina sequencing adapters. We pooled the final PCR product and sequenced using an Illumina NovaSeq X. Raw data are available in the National Center for Biotechnology Information (NCBI) Sequence Read Archive under the bioproject accession number <u>PRJNA1333493</u>.
**(iii)** **Data Processing and Statistical Analysis.** Raw NGS reads were quality filtered with fastqc and Trim Galore (https://github.com/FelixKrueger/TrimGalore)^47^. Our library inserts were designed such that the first 100 nucleotides are uniquely distinguishable even allowing up to 3 mismatches between query and subject. Sequencing reads were mapped to a reference of VSG phage library inserts using the bowtie aligner v1.2.3 (run parameters: –v 3 – m 1)^48^. The number of reads detected per phage clone were counted in sorted alignment files for each sample with HTSeq v.2.0.2 to create a read count matrix^49^. Fold change and significance testing per peptide was computed using EgdeR^50^ via the BEER R package (https://github.com/athchen/beer) which has been benchmarked for analysis of PhIP-seq data in a previous study^51^. The readout for PhIP-seq is a log2(fold change) value from EdgeR analysis of pairwise comparisons between each sample and the distribution generated from mock immunoprecipitation reactions with no serum. Significance is determined from these values if there is enrichment of reads belonging to a peptide over those detected as background signal in the mock distribution. Significantly enriched peptides required read counts ≥100 and Benjamini-Hochberg adjusted p-values ≤ 0.05 in at least 2 of the 3 technical replicates run per serum sample on each plate. We ran 8 beads-only mocks per plate, and all samples were compared against the mocks which were run on the same plate. The fold change matrix contains the average fold change value over mocks for all technical sample replicates which passed library quality control checks of at minimum 10x detected phage library read depth and 70% of reads successfully aligned to reference. All negative fold change values were set to zero (0) and those specificities considered undetected in a sample.

All downstream analyses for significantly enriched peptides were implemented in R v. 4.3.2 and plots created using the Tidyverse package suite. Matrices of Bray-Curtis dissimilarity or Jaccard distance were built with the Vegan R package (https://vegandevs.github.io/vegan/) for multi-dimensional scaling (MDS) plots. Bray-Curtis is often used in ecological studies to assess species diversity between communities by accounting for both the presence of a species and its abundance. In our case, we consider the abundance of a peptide to be its signal amplitude in Log_2_(fold change) of serum versus mocks. Thus, the calculation is unaffected by differences in sampling size. Heatmaps and associated hierarchical clustering of PhIP-seq signal amplitudes was done with the pheatmap R package. Scripts used for data analysis and figure generation are available at https://github.com/mugnierlab/So2025.

### Sequence Comparisons Between VSGs

We assessed the relatedness between peptide sequences, the basis for analysis of redundancy within the peptide library as well as for identification of epitopes, by adapting the network approach described in AVARDA (https://github.com/drmonaco/AVARDA) which provides a conservative estimate of the number of non-redundant groups of peptides^28^. Networks were built using igraph R software^52^ based on relations determined by pairwise blastp^53^ v.2.9.0 alignments between 15aa sequences tiled across all library peptides with a sliding window of 3aa and full length peptide tiles. A custom python script was used to break query sequences up into 15-mers and perform a blastp alignment to a reference of peptide tiles or assembled VSGs. Connections were drawn between any two peptides producing a blast hit using default parameters under –task blastp-short. This differs slightly from AVARDA which is based on shared 7-mer identical motifs, whereas this approach allows for identification of discontinuous sequence homology, which is often observed with antibody epitopes. From the blastp output, we built an edge list defining pairs of peptides with shared sequence for use with igraph to analyze redundancy within the library and define epitopes.

**(i)** **Library redundancy analysis.** The pairwise blastp output used to build the edge list for igraph was also useful for identifying spans of homology between missing and detected peptide tiles. The amount of redundancy between peptide tiles could be calculated by evaluating coverage of full-length peptides by the aligned15aa tiles. We determined how much of the sequence within a peptide tile was unique and how much could be found within other well<u>-</u> represented phage library tiles. A custom python script was used to perform this analysis. We revisit this approach throughout our analysis for estimating the amount of unique sequences represented by our phage pool, predicting epitopes, and for comparing phage library inserts to novel VSGs assembled from sequenced field isolates.

### Determining Immunogenic Regions of VSG Protein

**(i)** **Predicting and Mapping Putative Epitopes.** Peptides with redundant sequences often enriched together in the same serum sample. We assumed that peptides with highly similar sequences that co-enriched in the same patient were likely bound by the same antibodies, so any sequences shared between them could constitute epitopes. The edge list described above was used to generate network plots and perform unsupervised clustering of enriched peptides with shared homology for each individual patient using the cluster_leading_eigen() algorithm within igraph. Next, we determined the putative epitopes represented within each cluster as follows:
  - Peptide tiles that failed to cluster were assumed to contain unique antigenic sequence and the whole 90 amino acid tile was defined as the epitope.
  - We identified and extracted common sequences shared by small clusters consisting of 5 or fewer peptide tiles from the pairwise alignment used to generate the network plots.
  - Significantly enriched sequence motifs were identified in clusters of more than 5 peptide tiles using the MEME tool (https://meme-suite.org/meme/tools/meme) run under default parameters. Epitopes were then stringently mapped with ProteoMapper (http://www.tppms.org/pm/) to a collection of non-redundant full-length VSG sequences with known NTD types. We were concerned about highly similar VSGs skewing results, so VSGs with > 95% identity were collapsed to a single representative using cd-hit (parameters: –c 0.95 –G 0 –g 1 –aS 0.8 –n 5 – d 0 –M 0 –p 1). To find out which region of the VSG tends to be the most immunogenic, coordinates of all mapped epitopes per VSG were summarized with iranges R package. Epitope mapping coordinates for each VSG were normalized based on their position relative to the boundary between the NTD and CTD.
**(ii)** **Modeling Tertiary VSG Structure.** We sought to predict the structures of VSG NTDs. There are few reference structures that feature the whole VSG protein, and the CTD is inherently very disordered, so its structure cannot be predicted with high confidence at this time. Previous studies have shown AlphaFold2 to produce robust structural predictions for the NTD so we followed their approach^10^. In summary, we preprocessed VSG sequences to isolate the NTD using our Hidden Markov Model-based classification script. Signal peptides are not present in mature VSG proteins, so we predicted and removed these sequences using the SignalP 6.0 command line application (https://github.com/fteufel/signalp-6.0)^54^. Monomer structures of processed NTD sequences were predicted using the “colabfold_batch” application (parameters: –-templates –-amber –-num_recycle 3 –-use_gpu_relax) from LocalColabFold (https://github.com/YoshitakaMo/localcolabfold)^55,56^. Molecular graphics and analyses for predicted VSG NTD structures were performed with UCSF ChimeraX^57^ developed by the Resource for Biocomputing, Visualization, and Informatics at the University of California, San Francisco, with support from National Institutes of Health R01-GM129325 and the Office of Cyber Infrastructure and Computational Biology, National Institute of Allergy and Infectious Diseases.

### VSG Repertoire Characterization of Field Isolates

We assembled full-length VSG ORFs from whole genome Illumina short-read datasets. In total we analyzed 100 strains of *T. b. gambiense* type 1, 6 strains of *T. b. gambiense* type 2, 9 strains of *T. b. rhodesiense*, and 7 strains of *T. b. brucei*. Data were obtained from ENA, SRA, or Data Dryad repositories. Accession numbers and known metadata for each strain can be found in the Github repository associated with this publication. After running into complications assembling full contigs from the data, we opted instead to directly assemble ORFs and identify VSGs using a modified version of the VSG-seq pipeline available at https://github.com/mugnierlab/VSGSeqPipeline. Reads are assembled *de novo* using Trinity (version 5.26.2) and ORFs are identified within the VSG-seq script calling with a minimum protein length cutoff of 300 amino acids. VSGs are identified by blastn alignment of contigs to a reference of Lister 427 and EATRO1125 VSGs (E value < 1e-10) while filtering out known contaminating non-VSG sequences. Due to the unique biology of Trypanosomes, namely polycistronic transcription and complete lack of introns, we can assemble ORFs from genomic sequencing reads using Trinity which is designed for transcriptomic assembly.

### Statistical Machine Learning Classification Analysis

We created a matrix containing the average enrichment log2(fold change) value of technical replicates for all samples and detected peptides. There were over 30000 seropositive peptides between all samples which could have been used as predictors. The full datasets included all endemic negative controls and either gHAT patient or rHAT patient samples. We preprocessed the data to select smaller subsets of peptides prior to fitting any models. Initially we used Wilcoxon rank sum statistical testing. For rHAT classification we selected all peptides that met a Benjamini-Hochberg adjusted p-value < 0.2, yielding a short list of 24 peptides. Thousands of peptides met these thresholds in the gHAT dataset, so feature selection was performed using recursive feature elimination (RFE) with the Caret package in R. Optimal peptides selected with either RFE or significance testing were then used to fit models using various machine learning methods within caret. Without access to additional external data to use as a test set to evaluate the model, we instead set aside a portion of our PhIP-seq results which would not be used for model fitting. Training data consisted of 80% of the complete dataset randomly partitioned but maintaining the same proportions of each sample type, and remaining samples were the testing data for evaluating optimized models. We used caret to fit models using logistic regression (glm), linear discriminant analysis (LDA), boosting, and multivariate adaptive regression splines (MARS) which are various machine learning methods available within the package. Caret performed internal repeated k-fold cross validation (k = 5, repeats = 10) and found optimal models based on receiver operating characteristic (ROC) statistics. We then evaluated output models by using them to make predictions for the testing dataset and chose the best performing models for classifying gHAT (sensitivity > 0.95, test error < 0.0025) and rHAT (sensitivity > 0.70, test error < 0.15) patients. Candidate diagnostic peptides were identified by computing variable importance for the top models, another functionality of the Caret software package.

## Acknowledgements

We are very grateful to the patients and participants without whom this work would not have been possible. We thank Annette MacLeod for consulting with us about field isolates of *T. brucei* subspecies. We thank H. Benjamin Larman for guidance regarding phage display library design and implementing the PhIP-seq methodology. This work was carried out at the Advanced Research Computing at Hopkins (ARCH) core facility (rockfish.jhu.edu), which is supported by the NSF grant number OAC1920103. JS was supported by NIH 5T32AI007417.

